# Pairwise genetic interactions modulate lipid plasma levels and cellular uptake

**DOI:** 10.1101/2020.10.29.360818

**Authors:** Magdalena Zimon, Yunfeng Huang, Anthi Trasta, Jimmy Z. Liu, Chia-Yen Chen, Aliaksandr Halavatyi, Peter Blattmann, Bernd Klaus, Christopher D. Whelan, David Sexton, Sally John, Wolfgang Huber, Ellen A. Tsai, Rainer Pepperkok, Heiko Runz

## Abstract

Genetic interactions (GIs), the joint impact of different genes or variants on a phenotype, are foundational to the genetic architecture of complex traits. However, identifying GIs through human genetics is challenging since it necessitates very large population sizes, while findings from model systems not always translate to humans. Here, we combined exome-sequencing and genotyping in the UK Biobank with combinatorial RNA-interference (coRNAi) screening to systematically test for pairwise GIs between 30 lipid GWAS genes. Gene-based protein-truncating variant (PTV) burden analyses from 240,970 exomes revealed additive GIs for *APOB* with *PCSK9* and *LPL*, respectively. Both, genetics and coRNAi identified additive GIs for 12 additional gene pairs. Overlapping non-additive GIs were detected only for *TOMM40* at the *APOE* locus with *SORT1* and *NCAN*. Our study identifies distinct gene pairs that modulate both, plasma and cellular lipid levels via additive and non-additive effects and nominates drug target pairs for improved lipid-lowering combination therapies.

## INTRODUCTION

Genome-wide association studies (GWAS) have firmly established that changes in blood lipids and the risk of coronary artery disease (CAD) are heritable. Hundreds of genetic loci have been identified that reach genome-wide significant associations with plasma levels of low-density lipoprotein cholesterol (LDL), high-density lipoprotein cholesterol (HDL), triglycerides (TG), total cholesterol (TC) and CAD^1–4^. In rare instances, susceptibility to altered blood lipids can be attributed to mutations in individual genes such as *LDLR*, *PCSK9* or *APOB* that lead to familial forms of disease. For the vast majority of dyslipidemic individuals, however, no single-gene mutation can be identified. Instead, recent evidence suggests that in these cases inherited susceptibility is caused by a cumulative effect of numerous common alleles within and across GWAS loci. Individually, such common alleles have only a minor effect, but when summarized in polygenic scores they can modify a phenotype to a similar extent as single high-impact mutations^5^, or further magnify the penetrance of individual mutations causing Mendelian disease^6^. The biological mechanisms behind the cumulative effect of risk alleles in different genes remain unclear.

While the refined understanding of the polygenic nature of complex disease is starting to show promise for improved risk prediction and treatment decisions^7,8^, it has made it increasingly difficult to decide which individual genes could be the most suitable targets for developing new drugs. Drug development is traditionally focused on discrete targets with well-understood biology. For certain diseases, an additive therapeutic benefit has been demonstrated through combination therapies that simultaneously modulate two or more targets at once. For instance, combinations of statins, inhibitors of HMG-CoA-reductase (*HMGCR*), with distinct other cholesterol-lowering medications including *NPC1L1*, *PCSK9* and *APOB* inhibitors have been demonstrated to lower LDL levels and CAD-risk further than statin-treatment alone^9,10^. Despite such successes, systematic strategies to predict that joint modulation of drug target pairs in combination therapies will show benefit beyond standard of care have yet to be explored.

Genetic support for a drug target increases the probability that a medicine directed against the respective target will succeed by several fold^11,12^. We thus hypothesized that genetics might also assist in nominating drug target pairs that, when addressed jointly, will have a higher probability to reach a desired therapeutic benefit. A particular attractive approach to prioritize optimal target pairs would be to leverage synergistic gene-gene interactions, where genetic variants in two disease risk genes induce a phenotype that is more pronounced than what would be expected from each of the variants’ individual effects. Non-additive genetic interactions (naGIs), or epistasis, have been extensively studied in model organisms and cell models with the aim to identify functional relationships among genes and gene products^13,14^. In humans, however, the contribution of naGIs to the architecture of complex traits has been controversial. While there is increasing evidence for modifier genes that modulate Mendelian phenotypes in non-additive manners^15^, most of the variance of complex traits appears to be explained by genes acting additively within or between loci (additive GIs, or aGIs)^16^.

Here we systematically test for pairwise GIs regulating blood lipid levels by studying interactions between 30 genes prioritized based on known lipid-regulatory functions from GWAS loci using three complementary tools: protein-truncating variants (PTVs) identified through exome sequencing in the UK Biobank; reported GWAS lead SNPs genotyped or imputed in the UK Biobank; and combinatorial RNA-interference (coRNAi) screening measuring LDL-uptake into cultured cells. Our combined genetics and functional genomics approach establishes pairwise additive and non-additive GIs as foundational elements in controlling blood lipid levels and highlights distinct gene pairs as promising targets for lipid lowering combination therapies.

## RESULTS

### Study outline

To explore pairwise interactions between genes in GWAS loci and how these impact plasma lipid levels and LDL-uptake into cultured cells, we followed three parallel approaches: First, we extracted protein-truncating variants (PTVs) from whole exome sequencing data of 200,654 participants of the UK Biobank. Second, we utilized GWAS lead SNPs commonly used to construct polygenic risk scores from the full set of 378,033 unrelated participants of European ancestry in the UK Biobank. And third, we conducted systematic RNAi-based combinatorial knockdown experiments in cells (**Figure 1a**). We focussed our analyses on 30 high-confidence candidate genes from 18 genomic regions associated with blood lipid levels or the risk for CAD (**Table S1**). Twenty-eight of these genes had scored as functional regulators of LDL-uptake, cellular levels of free cholesterol, or LDL-receptor (LDLR) mRNA or protein levels in an earlier study where we had functionally analysed 133 genes at 56 lipid and CAD GWAS loci through RNAi-based knockdown experiments^17^. Causality for several of these genes to drive GWAS associations was further supported through systematic colocalization of plasma LDL GWAS lead SNPs with GTEx liver eQTLs^1^ (2 genes), cis-pQTL signals^18^ (3 genes) and independently reported biological evidence for lipid-relevant functions (15 genes) (**Table S2**). To identify pairwise GIs, we applied four linear regression models (modified from Axelsson et al., 2011^19^) to model the data. For each gene pair, both the additive genetic interaction effect (aGI) (model 3), which measures the sum of effects from each gene or variant individually, as well as the non-additive genetic interaction effect (naGI) (model 4), which measures the difference between the expected additive and the observed combined effect, were calculated, with a naGI being either *synergistic* or *buffering* (**Figure 1a** and Methods). Pairwise analyses were conducted for four plasma lipid parameters (LDL, HDL, TG, TC) and CAD as available from UK Biobank^20^ (see Methods).

**Figure 1.**
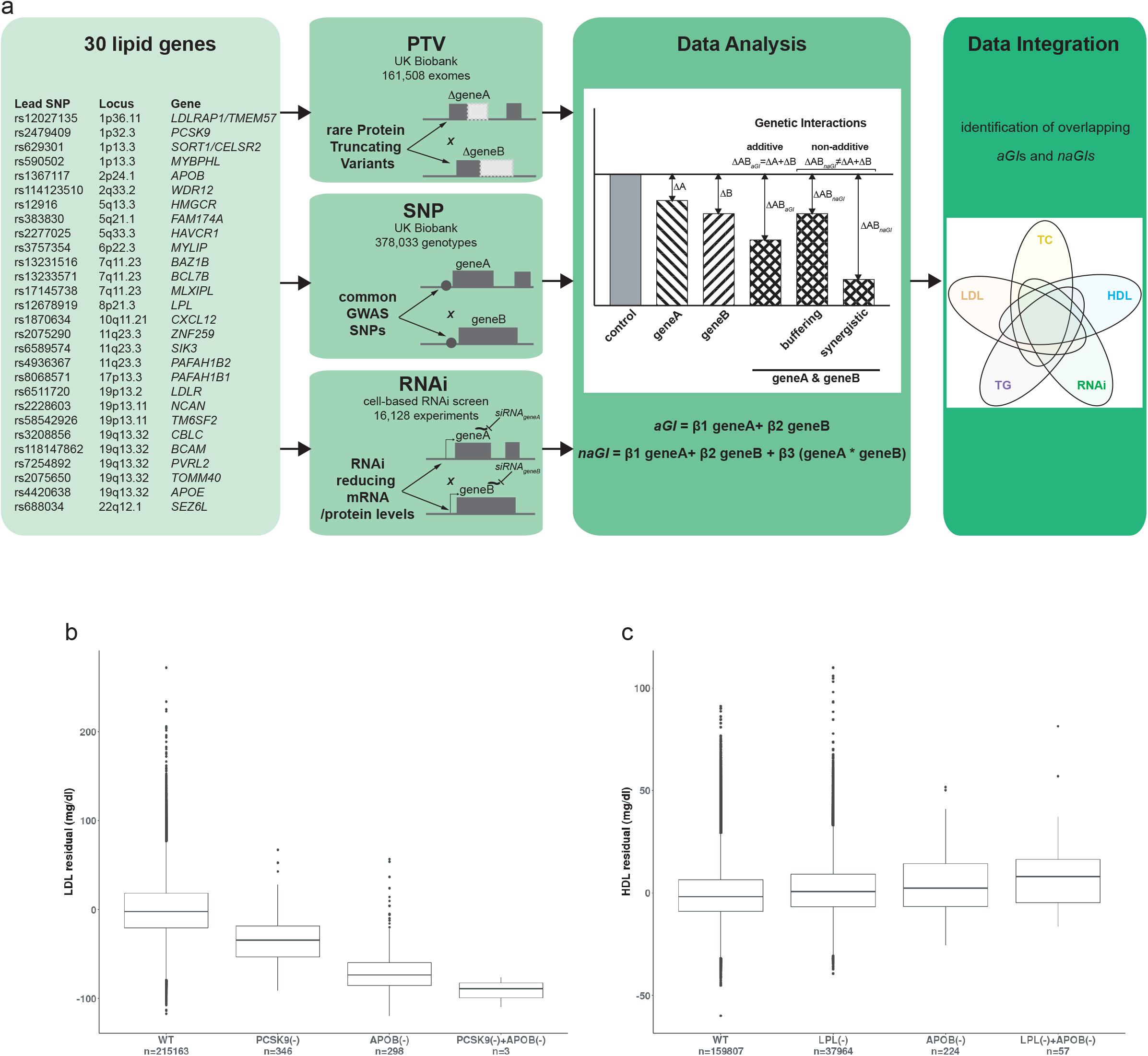
PTV-burden tests in UK Biobank establish additive GIs for *PCSK9*-*APOB* and *LPL*-*APOB*. **(a)** Workflow of the study. 30 high-confidence candidate genes for GI testing were chosen from 18 GWAS regions associated with blood lipid traits or CAD risk based on colocalization analyses with eQTL/pQTL signals and previously reported lipid-regulatory functions (see Methods). Pairwise GI analyses were conducted from three complementary datasets: protein-truncating variants (PTVs) from exome sequencing in the UK Biobank; lipid/CAD GWAS lead SNPs; and combinatorial RNAi (coRNAi) experiments in cells. Robust linear model fitting was used to identify additive (aGI) and non-additive (naGI) GIs, and genetic and functional data were integrated. **(b)** Gene-based PTV-burden analyses from 161,508 exomes identified an additive GI (aGI) for LDL (and TC; not shown) between *PSCK9* and *APOB*. (**c**) A suggestive non-additive GI (naGI) for HDL (and TG; not shown) between PTVs in *LPL* and *APOB* was validated as aGI for HDL, TG, and also LDL by replication analyses in an additional 79,462 UK Biobank exomes (Table S6). n, number of carriers. (-), predicted loss-of-function due to PTVs.

### PTV burden tests in UK Biobank reveal additive genetic interactions for *PCSK9*-*APOB* and *LPL*-*APOB*

We first studied pairwise modifier effects between the 30 candidate genes using high-impact protein-truncating variants (PTVs). PTVs are expected to cause loss-of-function and compared to other types of mutations are rare at the population level due to purifying selection^21,22^. We sequenced the exomes of 200,654 UK Biobank participants, annotated PTVs using Variant Effect Predictor v96^23^ and the LOFTEE plugin^21^, and identified 462,762 high-confidence PTVs in the canonical transcripts of 18,869 genes. Within the 30 lipid GWAS genes, we detected a total of 755 unique rare PTVs (**Table S3**). For instance, we discovered 29 different PTVs in *LDLR*, 47 in *PCSK9* and 102 in *APOB*. Most PTVs in these three genes were associated with strongly abnormal plasma LDL levels in heterozygote carriers consistent with Familial Hypercholesterolemia, although only 32 of the PTVs were annotated as pathogenic or likely pathogenic in ClinVar^24^.

Gene-based PTV-burden association analyses were conducted in a cohort of 161,508 unrelated UK Biobank participants of European ancestry. Single-gene PTV-burden testing identified three genes that were significantly associated (Bonferroni-corrected p<0.05) with both LDL and TC (*APOB*, *PCSK9*, *LDLR*), two with HDL (*LPL, APOB*) and two with TG (*LPL*, *APOB*), respectively (**Table S4**). Loss-of-function of these genes had already been identified earlier as associated with the respective lipid traits at the population level^2^. Next, we next expanded from these single gene PTV-burden analyses to study pairwise PTV-based GIs, which could be tested for 42 of the 435 theoretically possible gene combinations (**Table S5** and Methods). For the two gene pairs that met our stringent criteria to be classified as genetic interactions from this analysis, *PCSK9*-*APOB* and *LPL*-*APOB*, we conducted replication analyses in an additional 79,462 UK Biobank exomes, bringing the total sample size available for PTV-based GI testing to 240,970 individuals (**Table S6**). *PCSK9*-*APOB* showed an aGI for both, LDL and TC, reflecting that joint loss-of-function of both genes reduces these two lipid measures more than if only one of the two genes is truncated. For instance, PTVs in *PCSK9* and *APOB* individually reduced mean plasma LDL by 34.21 mg/dl and 69.42 mg/dl relative to individuals without PTVs in these genes, consistent with previous reports^25–27^. However, the three UK Biobank participants who carried both, *PCSK9* and *APOB* PTVs, showed on average a further reduction in plasma LDL by 40.01 mg/dl compared to individuals with PTVs in only one of the two genes, and by 90.45 mg/dl compared to individuals with no PTV in either of the two genes (**Figure 1b**), suggesting considerable additional protection from CAD. Additive GIs were further identified between *LPL* and *APOB* for HDL and TG. Individuals who carried PTVs in both, *LPL* and *APOB*, showed consistently higher HDL and TG levels than individuals with no PTVs, or PTVs in only one gene (**Figure 1c**). No naGIs were identified through PTV-based burden tests in up to 240,970 exomes. Our results are consistent with the prediction that for rare variant-based burden analyses very large sample sizes are necessary to robustly detect GIs in the human population^16,25^.

### Pairwise genetic interactions between GWAS loci modulate plasma lipid levels

We next tested for GIs using 28 lipid/CAD GWAS lead SNPs representing the 30 loci in 378,033 unrelated individuals of European ancestry in the UK Biobank^20^. Of a total of 1,890 pairwise SNP-SNP interactions tested, 195, 98, 124, 238 and 10 aGIs were identified for LDL, HDL, TG, TC and CAD, respectively (**Figure 2a-e**; **Table S7**). Interestingly, SNP-based analyses also suggested pairwise effects between GWAS loci that deviated from an additive model and were classified as naGIs. Specifically, we detected ten naGIs for LDL, one for HDL, six for TG, and nine for TC (**Table 1**). No naGI was detected for CAD. The strongest driver of interactions came from the 19q13.32 locus encompassing the *CBLC/BCAM/PVRL2/TOMM40*/*APOE* gene cluster that was contributing to 19 of the 26 naGIs identified across all traits. Fourteen naGIs were between lead SNPs from within the same GWAS region (“*cis*-naGI”, e.g., *NCAN*-*TM6SF2*, *BCAM-APOE*, *ZNF259-SIK3*) with nine of them being suggestive *cis*-effects of rs4420638 near *APOE*. However, naGIs were also identified between loci on different chromosomes (“*trans*-naGIs”), such as between *ZNF259* and *APOE,* or S*ORT1*/*CELSR2* and *TOMM40* for LDL and TC, or between *LPL* and *ZNF259*, or *LPL* and *SIK3* for TG. Overall, our data support the hypothesis that aGIs between GWAS loci are pervasive and individually small, yet if summed up across many loci in polygenic scores modulate complex traits^5^. Conversely, naGIs are considerably less prevalent, with the *APOE* locus being a potential contributor to naGIs for lipid traits.

**Figure 2.**
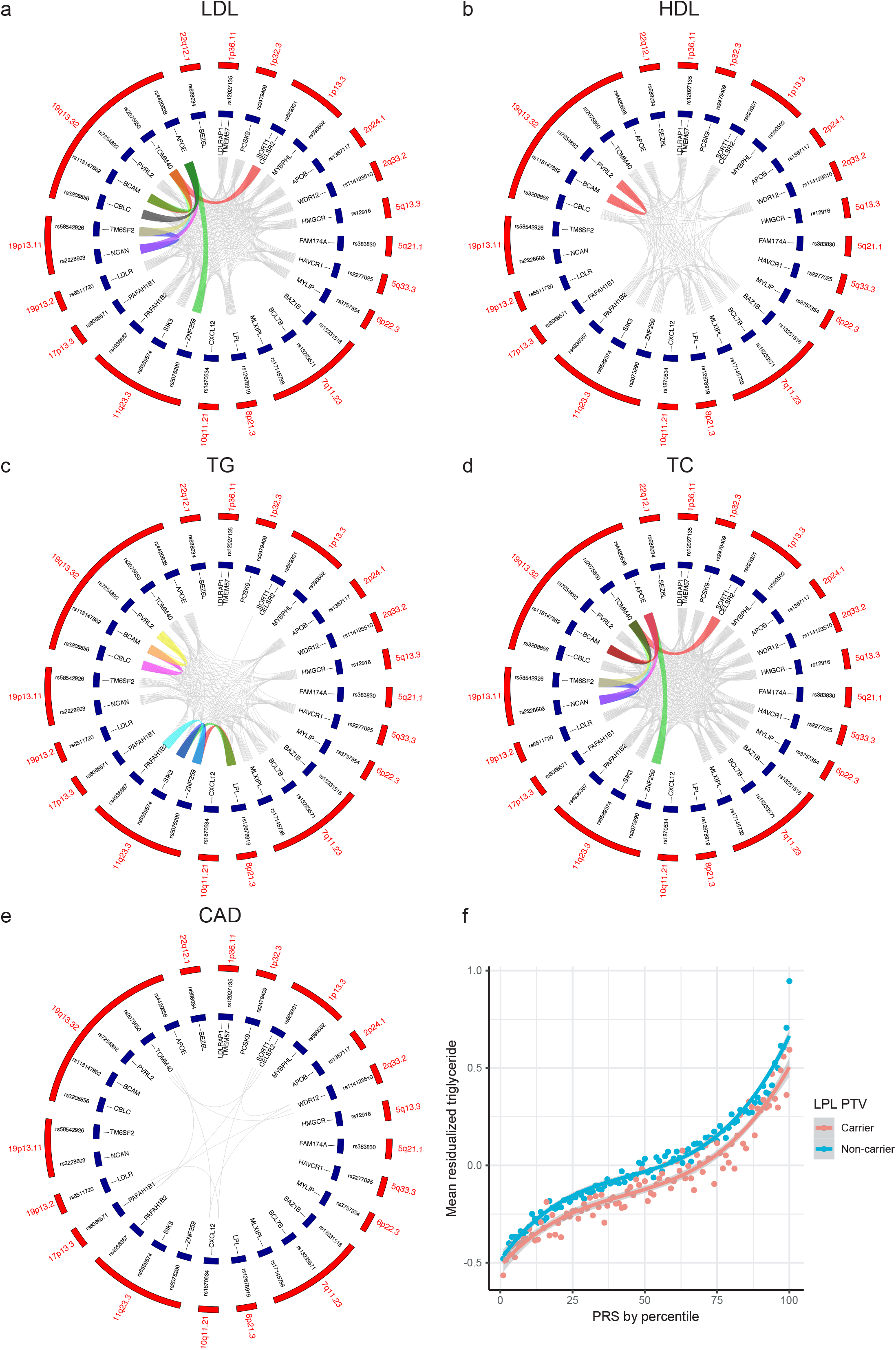
Pairwise GIs between lipid and CAD GWAS lead SNPs in 387,033 UK Biobank participants. **(a-e)** Circos plots showing aGIs (grey) and naGIs (colored) between GWAS lead SNPs (blue) at the 28 selected lipid/CAD loci (red) for the four tested lipid species and CAD. **(f)** Tests for GIs between polygenic risk scores for the four lipid species and PTV-burden for each of the 30 lipid genes identified a naGI between PTV-burden in *LPL* and the PRS for TG. PRS distribution (mean±SD) for *LPL*-PTV carriers (pink) and non-carriers (blue) are plotted against mean normalized residual TG levels. Each dot reflects mean TG levels at a respective percentile.

**Table 1.** Non-additive GIs from pairwise PTV-burden and GWAS lead SNP-based GI testing in UK Biobank.

| Trait | Gene 1 | Gene 2 | BIC_Best<br>_Model | Lowest<br>ΔBIC | incl.<br>19q13.32 | cis<br>GI | trans<br>GI | MAF<br>(SNP1) \|<br>N_PTV1<br>carriers | MAF<br>(SNP2) \|<br>N_PTV2<br>carriers |
| --- | --- | --- | --- | --- | --- | --- | --- | --- | --- |
| **LDL SNP-SNP GIs:** |  |  |  |  |  |  |  |  |  |
| LDL | NCAN (rs2228603) | TM6SF2 (rs58542926) | 4 | 85.34 |  | + |  | 0.076 | 0.076 |
| LDL | TM6SF2 (rs58542926) | APOE (rs4420638) | 4 | 63.34 | + | (+) |  | 0.076 | 0.191 |
| LDL | TM6SF2 (rs58542926) | TOMM40 (rs2075650) | 4 | 41.95 | + | (+) |  | 0.076 | 0.147 |
| LDL | BCAM (rs118147862) | APOE (rs4420638) | 4 | 34.25 | + | + |  | 0.046 | 0.191 |
| LDL | NCAN (rs2228603) | APOE (rs4420638) | 4 | 32.61 | + | (+) |  | 0.076 | 0.191 |
| LDL | NCAN (rs2228603) | TOMM40 (rs2075650) | 4 | 30.02 | + | (+) |  | 0.076 | 0.147 |
| LDL | BCAM (rs118147862) | TOMM40 (rs2075650) | 4 | 28.81 | + | + |  | 0.046 | 0.147 |
| LDL | ZNF259 (rs2075290) | APOE (rs4420638) | 4 | 4.68 | + |  | + | 0.068 | 0.191 |
| LDL | CBLC (rs3208856) | APOE (rs4420638) | 4 | 3.34 | + | + |  | 0.036 | 0.191 |
| LDL | SORT1/CELSR2 (rs629301) | TOMM40 (rs2075650) | 4 | 0.57 | + |  | + | 0.222 | 0.147 |
| **HDL SNP-SNP GIs:** |  |  |  |  |  |  |  |  |  |
| HDL | BCAM (rs118147862) | PVRL2 (rs7254892) | 4 | 1.47 |  | + |  | 0.046 | 0.031 |
| **TG SNP-SNP GIs:** |  |  |  |  |  |  |  |  |  |
| TG | ZNF259 (rs2075290) | SIK3 (rs6589574) | 4 | 31.81 |  | + |  | 0.068 | 0.084 |
| TG | BCAM (rs118147862) | PVRL2 (rs7254892) | 4 | 21.41 | + | + |  | 0.046 | 0.031 |
| TG | CBLC (rs3208856) | BCAM (rs118147862) | 4 | 20.45 | + | + |  | 0.036 | 0.046 |
| TG | ZNF259 (rs2075290) | PAFAH1B2 (rs4936367) | 4 | 17.82 |  | + |  | 0.068 | 0.1 |
| TG | LPL (rs12678919) | ZNF259 (rs2075290) | 4 | 13.81 |  |  | + | 0.098 | 0.068 |
| TG | LPL (rs12678919) | SIK3 (rs6589574) | 4 | 3.22 |  |  | + | 0.098 | 0.084 |
| **TC SNP-SNP GIs:** |  |  |  |  |  |  |  |  |  |
| TC | NCAN (rs2228603) | TM6SF2 (rs58542926) | 4 | 74.81 |  | + |  | 0.076 | 0.076 |
| TC | TM6SF2 (rs58542926) | APOE (rs4420638) | 4 | 53.33 | + | (+) |  | 0.076 | 0.191 |
| TC | TM6SF2 (rs58542926) | TOMM40 (rs2075650) | 4 | 38.17 | + | (+) |  | 0.076 | 0.147 |
| TC | NCAN (rs2228603) | APOE (rs4420638) | 4 | 30.93 | + | + |  | 0.076 | 0.191 |
| TC | NCAN (rs2228603) | TOMM40 (rs2075650) | 4 | 28.59 | + | + |  | 0.076 | 0.147 |
| TC | BCAM (rs118147862) | TOMM40 (rs2075650) | 4 | 11.24 | + | + |  | 0.046 | 0.147 |
| TC | BCAM (rs118147862) | APOE (rs4420638) | 4 | 9.05 | + | + |  | 0.046 | 0.191 |
| TC | ZNF259 (rs2075290) | APOE (rs4420638) | 4 | 2.40 | + |  | + | 0.147 | 0.191 |
| TC | SORT1/CELSR2 (rs629301) | TOMM40 (rs2075650) | 4 | 0.31 | + |  | + | 0.222 | 0.147 |
| **LDL PTV-SNP GIs:** |  |  |  |  |  |  |  |  |  |
| LDL | LDLR | PVRL2 (rs7254892) | 4 | 8.318 | + |  | + | 33 | 0.031 |
| **HDL PTV-SNP GIs:** |  |  |  |  |  |  |  |  |  |
| HDL | APOB | LPL (rs12678919) | 4 | 1.601 |  |  | + | 222 | 0.098 |
| **TC PTV-SNP GIs:** |  |  |  |  |  |  |  |  |  |
| TC | LDLR | PVRL2 (rs7254892) | 4 | 18.014 | + |  | + | 33 | 0.031 |
| TC | LDLR | SIK3 (rs6589574) | 4 | 2.961 |  |  | + | 33 | 0.084 |
| TC | LDLR | PAFAH1B2 (rs4936367) | 4 | 0.405 |  |  | + | 33 | 0.1 |
| **TG PTV-SNP GIs** |  |  |  |  |  |  |  |  |  |
| TG | LPL | SIK3 (rs6589574) | 4 | 8.4 |  |  | + | 31011 | 0.084 |
| TG | BAZ1B | PAFAH1B2 (rs4936367) | 4 | 5.825 |  |  | + | 25 | 0.1 |
| TG | LPL | ZNF259 (rs2075290) | 4 | 3.894 |  |  | + | 31011 | 0.147 |
| TG | BAZ1B | NCAN (rs2228603) | 4 | 2.845 |  |  | + | 25 | 0.076 |
| TG | LPL | PAFAH1B2 (rs4936367) | 4 | 1.648 |  |  | + | 31011 | 0.1 |
| TG | BAZ1B | TM6SF2 (rs58542926) | 4 | 1.522 |  |  | + | 25 | 0.076 |

Non-additive genetic interactions (naGIs) identified through GWAS lead SNP- and PTV-SNP-based GI analyses in the UK Biobank as described in Methods. BIC, Bayesian Information Criterion. A lowest ΔBIC of "4" indicates interaction model is most compatible with a non-additive interaction effect. MAF (minor allele frequency) estimates and numbers of rare protein-truncating variant (PTV) carriers are based on genotypes from 387,033 and exomes, respectively, from 161,508 unrelated UK Biobank participants of European ancestry. (+) indicates possible cis-effects of rs4420638 in APOE on neighbouring genes on Chr.19q13.32. Trans GI indicates genes contributing to pairwise naGIs are located on different chromosomes. \* represents data based on the replication study in additional 79,462 UK Biobank participants.

### Genetic interactions between gene-based PTV-burden and GWAS loci or polygenic scores

Next, we queried for GIs between different types of genetic variation. Pairwise interaction testing between gene-based PTV-burden and GWAS lead SNPs identified one naGI for LDL (*LDLR*_PTV_-*PVRL2*_SNP_), one for HDL (*APOB*_PTV_-*LPL*_SNP_), three for TC (*LDLR*_PTV_-*PVRL2*_SNP_, *LDLR*_PTV_-*SIK3*_SNP_, *LDLR*_PTV_-*PAFAH1B2*_SNP_), and six for TG (*LPL*_PTV_-*SIK3*_SNP_, *LPL*_PTV_-*ZNF259*_SNP_, *LPL*_PTV_-*PAFAH1B2*_SNP_, *BAZ1B*_PTV_-*NCAN*_SNP_, *BAZ1B*_PTV_-*TM6SF2*_SNP_, *BAZ1B*_PTV_-*PAFAH1B2*_SNP_) (**Table S8**). Moreover, 56, 26, 54 and 31 aGIs were identified for LDL, HDL, TC and TG, respectively. These results are consistent with the genetic architecture regulating plasma lipids being continuous between high-impact rare and low-impact common alleles^4^.

A recent study^6^ proposed that the penetrance of Mendelian disease, including FH, can be substantially modulated by interactions between the respective mutant gene with common variants (minor allele frequency >0.01) of individually small effect size subsumed in polygenic risk scores (PRS). We created PRS for the four lipid species using PRS-CS^26^ (and Methods) and tested for GIs between PRS and PTV-burden for each of the 30 genes. Of all combinations tested, only PTV-burden in *LPL*, mostly driven by the frequent p.S447Ter variant, showed evidence for a naGI with the PRS for TG (p<1.13×10^−15^; beta=-0.04) (**Figure 2f**; **Table S9**). This supports the hypothesis that a high polygenic risk for elevated TG can be mitigated by a concomitant stop-gain mutation in LPL. Additionally, 10 aGIs were identified between *APOB*_PTV_ with PRS for all four lipid species, *LDLR*_PTV_ and *PCSK9*_PTV_ with PRS for LDL and TC, and *LPL*_PTV_ with PRS for LDL and HDL.

### RNAi identifies pairwise functional gene interactions modulating cellular LDL-uptake

To gain insights into the functional consequences of GIs, we complemented our genetic analyses with systematic experiments in cells using combinatorial RNAi (coRNAi) (**Figure 3a** and Methods). We applied solid-phase reverse transfection to simultaneously knock down candidate gene pairs in cultured HeLa cells, which we have previously shown to reliably reflect various aspects of LDL biology and lipid homeostasis^17,27,28^. Each of the 30 lipid genes was profiled with a single siRNA that had previously been validated to significantly enhance or reduce cellular uptake of fluorescent-labelled LDL (DiI-LDL) or free cellular cholesterol levels, and/or to efficiently downregulate mRNA or protein levels of its respective target gene (**Table S2**)^17^. The impact of both, single and combinatorial gene knockdown on LDL-uptake per cell was measured and quantified from high-content microscopy images using automated image analysis routines as described (**Figure S1**)^27,28^. All pairwise knockdown combinations between the 30 lipid genes (435 gene pairs) were assayed in a total of 16,128 experiments (**Figure 3b**). Each combination was tested in at least seven biological replicates. Using BIC-model based robust linear regression fitting analogous to the genetic interaction analyses, we identified 18 aGIs and 33 naGIs to differentially impact cellular LDL-uptake (**Table S10**). A similar proportion of GIs was identified using robust linear model fitting and deriving p-values from the linear regression model term describing non-additive effects as an alternative statistical approach (see Methods). This identified 35 naGIs, with 31 naGIs overlapping between both analytical approaches (**Table S11**). The corresponding gene pairs were brought forward to independent liquid-phase based coRNAi replication experiments that validated 20 of these naGIs (**Table 2**, **Table S12, Figure S2**). Of the 20 validated naGIs identified through coRNAi, seven were classified (according to Horlbeck et al., 2018^14^) as *synergistic*, i.e., simultaneous knockdown of both genes magnified the effect size beyond expectations for an aGI; and thirteen naGIs were categorized as *buffering*, i.e., relative to an aGI the joint knockdown mitigated LDL-uptake into cells (**Figure 3c**). For instance, simultaneous knockdown of *HMGCR* and *APOB* enhanced cellular LDL-uptake beyond a mere additive effect expected from knockdown of either of the two genes, proposing a *synergistic* naGI (**Figure 3d**), that is most likely explained by a higher capacity of cells to bind and internalize LDL via increased availability of LDL-receptor at the cell surface (**Figure S3**). Conversely, knockdown of *LDLR* strongly inhibited, whereas partial knockdown of *LDLRAP1* increased cellular LDL-uptake under our experimental conditions. When silencing *LDLR* and *LDLRAP1* jointly, the reduction of LDL-uptake was less attenuated than expected under an additive model, suggesting a *buffering* naGI (**Figure 3e**). Interestingly, reduction of LDL-uptake upon knockdown of *LDLR* was magnified when *LDLR* was jointly silenced with *HAVCR1*, a suggested LDL scavenger receptor that might contribute to maintain the potential of LDLR-depleted cells to internalize LDL^29^ (**Figure 3f**). Noteworthy, among the remaining validated coRNAi naGIs, simultaneous silencing of *PCSK9* and *TMEM57*, as well as of *SIK3* and *PAFAH1B1* increased cellular LDL-uptake to a similar extent as the simultaneous knockdown of *HMGCR* and *APOB*, although silencing of these genes individually had a significant, yet only modest impact on cellular LDL-uptake. In summary, coRNAi identified aGIs and naGIs between established lipid-regulatory genes, but also proposed combinations of less well characterized genes as potentially important factors in maintaining cellular lipid levels.

**Table 2.** Pairwise GIs identified and validated through coRNAi to impact LDL-uptake into cells.

| GI Gene Pair |  | Interaction type | BIC-based GI testing |  | RLMF-based GI testing |  |  | Validation RNAi GI testing |  |  |
| --- | --- | --- | --- | --- | --- | --- | --- | --- | --- | --- |
| Gene1 | Gene2 | | Lowest ΔBIC | Best Model | Robust Zscore | Interaction Value | pVal(fdr) | Robust Zscore | Interaction Value | pVal (fdr) |
| *APOB* | *HMGCR* | Synergistic | 4.62 | 2 | 2.8 | 2.18 | 4.05E-03 | 3.33 | 0.97 | 1.53E-03 |
| *HAVCR1* | *LDLR* |  | 4.31 | 4 | -2.18 | -1.32 | 1.68E-02 | -2.24 | -1.24 | 1.81E-10 |
| *LDLR* | *NCAN* |  | 2.36 | 4 | -1.86 | -1.28 | 7.64E-03 | -1.23 | -2.22 | 2.20E-11 |
| *MYBPHL* | *SIK3* |  | 2.27 | 4 | 2.17 | 1.59 | 6.59E-03 | 2.3 | 0.96 | 7.91E-03 |
| *PAFAH1B1* | *SIK3* |  | 4.49 | 4 | 1.62 | 1.79 | 2.52E-03 | 3.3 | 2.44 | 8.37E-12 |
| *PCSK9* | *TMEM57* |  | 3.82 | 4 | 1.46 | 1.76 | 2.37E-03 | 3.21 | 2.44 | 3.74E-11 |
| *BCAM* | *LDLRAP1* | Buffering | 4.70 | 4 | -0.4 | -1.83 | 3.58E-04 | -0.05 | -0.7 | 5.20E-03 |
| *CELSR2* | *LPL* |  | 0.04 | 0/2 | -1.4 | -1.43 | 7.53E-03 | -0.11 | -0.86 | 8.70E-05 |
| *CXCL12* | *PAFAH1B1* |  | 5.47 | 4 | -2.17 | -1.78 | 3.69E-04 | 1.32 | -1.92 | 9.84E-13 |
| *HAVCR1* | *LDLRAP1* |  | 13.70 | 4 | -0.56 | -2.16 | 1.03E-05 | 0.39 | -0.73 | 5.40E-03 |
| *HAVCR1* | *MLXIPL* |  | 13.55 | 4 | -0.63 | -2.31 | 1.03E-05 | -0.12 | -1.2 | 2.06E-05 |
| *HAVCR1* | *SEZ6L* |  | 8.54 | 4 | -1.19 | -1.91 | 3.39E-04 | -0.67 | -0.77 | 2.90E-04 |
| *HAVCR1* | *SORT1* |  | 11.66 | 4 | -2.09 | -2.24 | 9.34E-05 | -0.63 | -0.58 | 1.98E-03 |
| *LDLR* | *LDLRAP1* |  | 10.06 | 4 | -2.44 | -1.86 | 9.34E-05 | -2.49 | -0.92 | 2.36E-10 |
| *LDLR* | *MLXIPL* |  | 5.79 | 4 | -2.03 | -1.49 | 2.97E-03 | -2.26 | -0.78 | 4.19E-11 |
| *LDLRAP1* | *SORT1* |  | 8.13 | 4 | -1.11 | -1.65 | 2.57E-03 | -1.09 | -0.95 | 3.89E-06 |
| *MLXIPL* | *TOMM40* |  | 18.73 | 4 | -2.14 | -2.67 | 1.03E-05 | -1.94 | -0.85 | 3.82E-08 |
| *NCAN* | *SEZ6L* |  | 5.08 | 4 | -0.59 | -1.6 | 5.54E-03 | -0.08 | -1.56 | 7.94E-12 |
| *NCAN\** | *TOMM40\** |  | 2.76 | 4 | -1.4 | -1.56 | 7.17E-03 | -1.49 | -1.34 | 1.75E-08 |
| *SORT1\** | *TOMM40\** |  | 5.80 | 0 | 0.77 | 1.84 | 3.90E-03 | -2.48 | 1.77 | 0.00E+00 |

**Figure 3.**
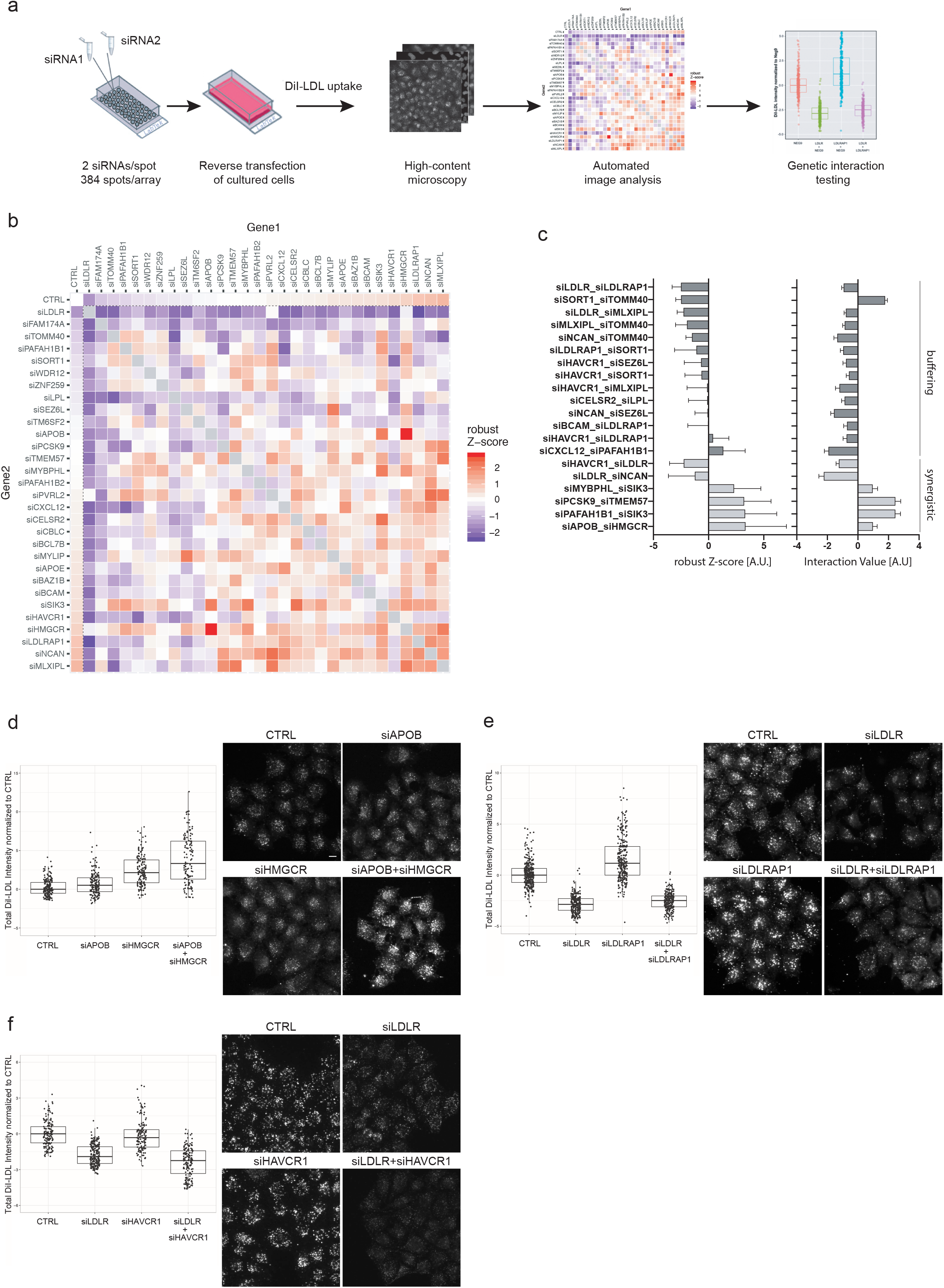
Combinatorial RNAi identifies pairwise GIs modulating cellular LDL-uptake. **(a)** coRNAi screen workflow. Customized cell microarrays were generated by pairwise spotting of siRNAs against two different candidate genes on 384 spots/array for solid-phase reverse siRNA transfection of cultured HeLa cells. Cells challenged to internalize fluorescent-labelled LDL (DiI-LDL) over a period of 20 min were imaged on a high-content microscope. Integrated fluorescence intensities for each cell individually were quantified by automated image analysis. Averaged signal intensities per gene pair were tested for GIs in multiple replica experiments per array. GIs suggested in the coRNAi screen as potentially non-additive were subsequently validated in customized experiments using fluid-phase transfection. **(b)** Heatmap visualizing median robust Z-score distribution upon coRNAi of 435 gene pairs assessed for their impact on cellular LDL-uptake. Red, increase. Blue, decrease. CTRL (top row and first column) reflects the relative impact on LDL-uptake when candidate genes were silenced individually (siRNA_*geneAorB*_ + negative control siRNA). **(c)**20 gene pairs validated as either buffering or synergistic naGIs on cellular LDL-uptake in independent replica experiments, sorted according to effect size. Interaction Value (right graph) depicts the directionality and difference of the combined effect versus single knockdown effects. **(d-f)** Selected examples of single gene (siRNA_*geneA*_ + negative control siRNA) and gene pair (siRNA_*geneA*_+siRNA_*geneB*_) siRNA knockdown effects on relative fluorescently-labelled LDL (DiI-LDL) cellular uptake. CTRL, control siRNA. Boxplots represent values between 25^th^ and 75^th^ percentile, whiskers indicate largest value within 1.5 times interquartile range above 75^th^ percentile. Median value is highlighted in the boxplot as a horizontal line. Dots represent robust Z-score values calculated for integrated DiI fluorescence intensities per cell (see Methods). Scale bar=10 μm.

### Integrated analysis highlights GIs supported by both human genetics and cellular function

In order to assess whether GIs identified through either PTV-based gene-burden tests, GWAS lead SNPs, or cell-based coRNAi overlapped, we integrated results from the three approaches (**Figure 4**; **Table S13**). *LDLR*-*SIK3* showed an aGI both in coRNAi and PTV-SNP analyses for LDL (**Figure 4a**). Both, coRNAi screening (ΔBIC 16.87, pVal(FDR)=1.18E-07) and PTV-SNP analyses for LDL and TC proposed a naGI between *LDLR* and *PVLR2* (**Figure 4b**), although this gene pair failed to score as naGI in the independent coRNAi validation experiments. Twelve of the 18 gene pairs nominated by coRNAi as aGIs also scored as aGIs in SNP-based interaction testing for LDL and TC, including *LDLR*-*SIK3*. Five aGIs involved *HMGCR* and four *LDLRAP1* (**Figure 4c**). Two gene pairs, *SORT1*-*TOMM40* and *NCAN*-*TOMM40,* scored as naGIs both in the SNP-based as well as the coRNAi-based interaction testing (**Figure 4d**), with *TOMM40* exerting a buffering naGI in either gene pair (**Figure 4e**) that could not be explained by an off-target effect of *TOMM40* siRNAs on *APOE* as an adjacent gene in the 19q13.32 GWAS locus (**Figure S4)**. In conclusion, integrating genetic with functional data validated 12 proposed aGIs and further substantiates a role of the *APOE* locus, and possibly *TOMM40*, as contributing to non-additive genetic interactions.

**Figure 4.**
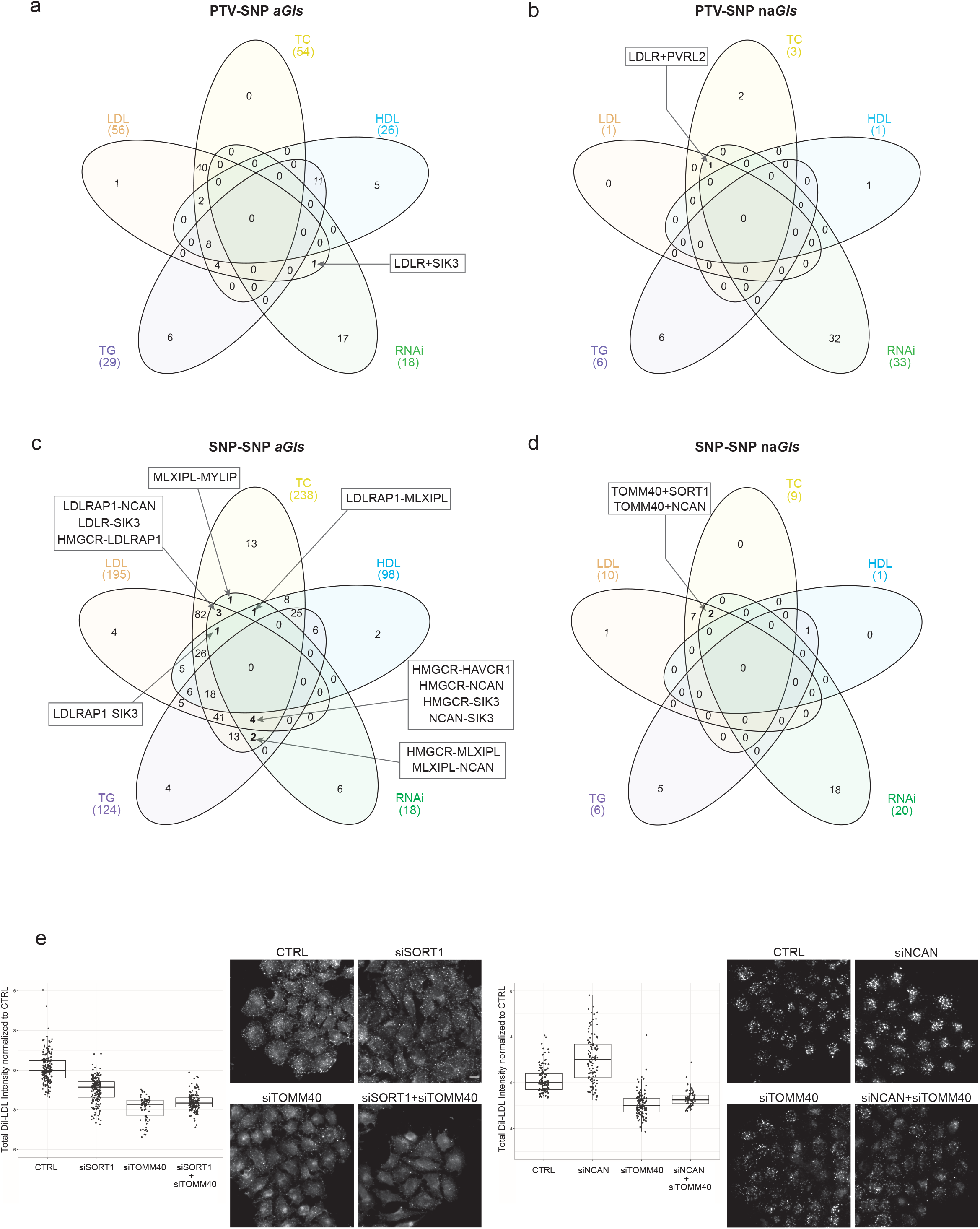
Integrative analysis identifies pairwise GIs supported by both, genetic and functional data. Overlap of GIs identified through genetic analyses and coRNAi. Highlighted are gene pairs identified through either PTV-SNP **(a, b)** or SNP-SNP **(c,d)** GI testing for which pairwise siRNA-knockdown showed corresponding effects on cellular LDL-uptake, validating these GIs as either aGI (a,c) or naGI (b,d). **(e)***TOMM40* as an example for which, consistent with SNP-SNP analyses, siRNA knockdown revealed buffering naGIs when jointly silenced with *SORT1* (left panel) or *NCAN* (right panel). Values on the graphs reflect robust Z-scores values calculated for total intensity of DiI-LDL per cell averaged per image (see Methods). Boxplots represent values between 25^th^ and 75^th^ percentile, whiskers indicate largest value within 1.5 times interquartile range above 75^th^ percentile. Median value is highlighted in the boxplot as a horizontal line. Dots represent robust Z-score values calculated for integrated DiI fluorescence intensities per cell (see Methods). Scale bar= 10 μm.

## DISCUSSION

Here, we apply whole-exome sequencing, genotyping and coRNAi to systematically test for pairwise GIs between 30 lipid-regulatory genes at lipid and CAD GWAS loci. GIs are considered to be central constituents of biological pathways and complex traits, contributors to human disease, and promising starting points for therapy development^13,15^. Mapping GIs, and particularly non-additive epistasis, however, has been challenging. GI studies require very large population sizes in order to obtain sufficient statistical power, so that the large number of potential interactions to be evaluated quickly leads to a prohibitive number of statistical tests^30^. Together with most GI studies to date being limited to just a single datatype, the relative contribution of GIs to variation in human complex traits has been controversial, and the relevance of epistasis potentially overestimated^16^.

In our study, we have tried to overcome several of these challenges through a systematic approach to GI testing that integrates genetic with functional data and relies on the UK Biobank, a population cohort linking genetic with phenotype data at an unprecedented scale^20^. To protect against statistical penalties from multiple hypothesis testing we focused on pairwise interaction analyses between 30 candidate genes nominated through GWAS that functional or genetic follow-up studies have proposed as likely causal to confer associations with lipid traits or CAD^17^. We assessed these genes for GIs across the allelic spectrum, from rare PTVs ascertained from the exomes of more than 240,000 individuals, to common GWAS lead SNPs. Genetic GI-testing was complemented by functionally knocking down gene pairs with siRNAs and determining the consequence on LDL internalization into cells.

Several of the GIs identified in our study can be expected to be high potential starting points for the development of advanced lipid-lowering combination therapies. Lowering LDL with statins is the first-line pharmacological strategy to treat or prevent CAD and ischaemic heart disease as its clinical manifestation. However, many patients do not reach their recommended goals of LDL-lowering through statins alone, or they are intolerant against statins. For these, combination therapies have become available that aim to lower atherogenic lipid levels further. A motivation for this is that every 1 mmol/l (39 mg/dl) reduction in blood LDL is associated with a 19% reduction in coronary mortality and a 21% reduction in major vascular events, supporting that, at least for secondary prevention, the lower blood LDL levels, the better^31^. Among the options that lower atherogenic blood lipids the most successfully are therapeutics against drug targets that when mutated cause familial hypercholesterolemia (FH), such as *NPC1L1*, the target of ezetimibe, or *PCSK9*^9^. Genetic analyses in extreme phenotypes have identified a small number of individuals with concomitant mutations in two distinct FH genes, such as *LDLR* and *APOB*^32,33^, *LDLR* and *LDLRAP1*^34,35^ or *APOB* and *PCSK9*^36^. However, due to the rarity of highly penetrant FH mutations such findings have thus far remained limited to individual families. Conversely, on a population level, a previous GI analysis based on common alleles from ~24,000 individuals ascertained for lipid traits reported 14 replicated GIs between lipid GWAS loci, most notably, like in our study, with SNPs at the *APOE* locus being a key contributor^37^. Additional support for the relevance of GIs for modulating lipid traits comes from a recent study that includes a subset of the UK Biobank exomes analysed here and proposes an interplay of genetic variation across the allelic spectrum^6^. Notably, that study reports that carriers of monogenic CAD risk variants show an up to 12.6-fold higher risk to manifest disease if they are in the highest quintile of the polygenic risk distribution.

Our analyses here propose distinct gene pairs that modulate plasma and cellular lipid levels via additive and non-additive GI effects. Among others, we identify GIs for several prominent cardiovascular risk genes that individually are established targets for lipid-lowering drugs. For instance, coRNAi proposed a synergistic, non-additive GI between *HMGCR*, the rate-limiting enzyme during cholesterol biosynthesis and target of statins, and *APOB* encoding apolipoprotein B, a critical constituent of LDL particles. Consistent with the known biological functions of these genes, joint knockdown increased levels of functional LDL-receptor on the cell surface and stimulated internalization of exogenous, fluorescent-labelled LDL. This observation is well in line with results from clinical trials showing that in patients with Familial Hypercholesterolemia and other hyperlipidemias a combination of statins with an antisense inhibitor of apolipoprotein B (mipomersen) efficiently reduces plasma LDL levels more strongly than high-intensity statin treatment alone^38–41^. Importantly, the additive GI identified from UK Biobank participants carrying PTVs in both, *APOB* and *PCSK9* suggests that similarly beneficial effects can be expected when APOB antisense therapies are applied in combination with PSCK9 inhibitors. Recently, inclisiran, an siRNA targeting PCSK9 in individuals on maximally tolerated statin doses^42^ led to a persistent, highly significant lowering of LDL in treated individuals relative to placebo in a phase 3 study^43^, introducing siRNAs as an attractive therapeutic modality for lipid-lowering therapies. Our results strongly propose that, on a population level, combination therapies inhibiting both *PCSK9* and *APOB* may lower LDL-C levels and CAD-risk even more substantially than drugs targeting only one of the two genes.

*APOB* PTV-burden was associated not only with LDL and TC, but also HDL and TG, and our PTV-based GI tests propose that joint disruption of *APOB* together with *LPL* reduces TG and increases HDL, most likely in an additive manner. *LPL* encodes for lipoprotein lipase which hydrolyzes TG from apolipoprotein B containing lipoproteins, releasing fatty acids^44^. PTV-burden in *LPL* is dominated by the stop-gain variant p.Ser447Ter (c.1421G>C; rs328) which in our exome-sequenced UK Biobank sub-cohort showed an allele frequency of 9.95%. This variant is known to cause gain-of LPL activity leading to a 0.8-fold reduced risk for ischaemic heart disease^45^, an effect that is likely to be further enhanced by concomitant reduction of apolipoprotein B. The p.Ser447Ter allele was also the main driver behind the only naGI detected between PTV-burden and polygenic risk for plasma lipids and conferred that in *LPL* PTV-carriers polygenic risk for TG is reduced, with presumably non-additive effects being the most pronounced in the upper percentile range of the PRS distribution.

A prominent driver of GIs in both our SNP- and coRNAi-based analyses was the 19q13.32 locus which includes *APOE* and apart from plasma lipids and CAD is associated with Alzheimer’s disease, longevity and macular degeneration among others^18^. Interestingly, our findings indicate that genes other than *APOE* at this locus might contribute to lipid GIs, which is consistent with our earlier findings that knock down of several genes at this locus independently modulate cellular LDL-uptake^17^. For instance, both SNP-based GI testing and coRNAi suggested buffering naGIs for *TOMM40* with *SORT1* and *NCAN*, respectively. Variants in *TOMM40* have been hypothesized to modify onset of Alzheimer’s disease independently of and in conjunction with *APOE*^45^. Our analyses suggest *TOMM40* might exert similar modifying effects on lipid phenotypes and CAD risk, which will need to be clarified in future studies. Another gene at the 19q13.32 locus is *PVRL2*, for which both coRNAi and SNP-PTV analyses proposed GIs with *LDLR*. As a vascular cell adhesion molecule, PVRL2 protein regulates transendothelial migration of leukocytes. PVRL2 levels in the atherosclerotic arterial wall correlate with plasma cholesterol in CAD patients and *Ldlr*-deficient mice and have been linked to the progression of atherosclerosis^46,47^. It is thus tempting to speculate that the extensive pleiotropy of the 19q13.32 locus can at least in part be explained through non-*APOE* related mechanisms^45^.

Both, genetic and functional analyses further revealed GIs between *HAVCR1*, *NCAN* and *SIK3* with *HMGCR*, nominating these poorly characterized genes to be explored as potentially attractive new targets for lipid-lowering therapies on top of statins.

Consistent with previous assumptions^16^, our results show that for regulating plasma lipid levels, additive GIs between gene or variant pairs are common, while non-additive epistasis is rare. Indeed, despite a sample size of over 240,000 exomes, our gene-based PTV-burden GI analyses did not find evidence for pairwise naGIs between lipid genes disrupted by PTVs. Further increasing sample sizes might help uncover non-additive effects, however, at least for lipid traits, their contribution to the overall variance appears to be small. This is consistent with the existence of evolutionary mechanisms that suppress epistatic interactions^13^. Since pairwise naGIs can be expected to be identified the most easily for genes that are disrupted sufficiently frequently in a population by PTVs of large-enough effect size, sequencing of consanguineous or bottlenecked populations might improve the detection rate of naGIs^22,48^. Interestingly, as observed also here, naGIs seem to be more easily detectable in cell and animal models, for instance through synthetic lethality mapping^14^.

Integration of population-scale genetics and functional coRNAi screening results yielded a total of twelve aGIs and three naGIs (one of them suggestive) that influence plasma and cellular lipid levels. Such validation via two systematic approaches substantially increases the confidence for committing to time and resource-intense follow-up analyses of such findings, e.g., when exploring the suitability of a gene pair to be jointly targeted in combination therapies. Interestingly, a significant number of GIs identified through genetics and coRNAi in our study do not yet overlap. This may be explained by several reasons: First, our functional analyses were limited to measuring LDL-uptake into cells, which reflects a relevant, yet only a partial aspect of the many possible mechanisms by which a gene can modulate plasma lipid levels. Second, siRNA-based gene knock down captures acute and rather severe functional effects, which may differ from the chronic and often compensated consequences upon lifelong modulation of a gene’s function through genetic variation. Third, despite the large number of samples used for genetics-based GI testing, the number of informative high-impact variants in the human germline may still be too discrete to comprehensively identify GIs. Regardless, the availability and rapid development of advanced high-throughput microscopy technology joint with the constantly increasing cohort sizes for genetic analyses will allow up-scaling of the approach taken here in future studies and with a high probability validate further GIs.

In conclusion, our study introduces and confirms a strategy to link large-scale genetic data from a population biobank with quantitative, cell-based coRNAi to map GIs that affect blood lipid levels and CAD, an approach that can be applied to other diseases and complex traits. Our unbiased analyses support that mechanisms exist through which multiple genes jointly help maintain blood lipid homeostasis. CAD and ischaemic heart disease remain a substantial global health burden, and doubling-down on lowering atherogenic plasma lipids remains one of the most promising therapeutic approaches. With the encouraging results from recent gene- and antisense-based clinical trials for CAD, our results help prioritize drug target pairs for the development of lipid-lowering combination therapies rooted in human genetics.

## ACKNOWLEDGEMENTS

This research has been conducted using the UK Biobank resource under application number 26041. We thank all the participants and researchers of UK Biobank for making these data open and accessible to the research community. AbbVie, Anylam Pharmaceuticals, AstraZeneca, Biogen, Bristol-Myers Squibb, Pfizer, Regeneron and Takeda are acknowledged for generation and initial quality control of the whole-exome sequencing data. We thank Eric Marshall, Yongsheng Huang and Frank Nothaft for infrastructure support for genetic data analyses. The EMBL Advanced Light Microscopy Facility is acknowledged for supporting high-content microscopic-based screening analyses. We are grateful to Brigitte Joggerst, Susanne Theiss and Miriam Reiss for excellent technical assistance. Support to the study came in part from the Transatlantic Networks of Excellence Program 10CVD03 from Fondation Leducq to HR and RP. MZ was supported by the EMBL EIPOD programme, AT and PB by the EMBL PhD programme.

## AUTHOR CONTRIBUTIONS

Conceptualization, H.R. and R.P.; Methodology and Investigation, M.Z., Y.H., A.T., C.-YC., R.P. and H.R.; Formal Analysis and Validation, M.Z., Y.H., A.T., C.-Y.C., J.L., A.H., B.K., E.T. and H.R.; Resources and Data Curation, J.L., P.B., C.W., D.S., S.J., E.T., R.P. and H.R.; Writing – Original Draft, M.Z., A.T. and H.R; Writing – Review and Editing, M.Z., Y.H., P.B., E.T., R.P. and H.R.; Supervision, S.J., W.H., E.T., R.P. and H.R.; Project Administration and Funding Acquisition, S.J., R.P. and H.R.

## DECLARATION OF INTERESTS

Y.H., C.-Y.C., J.L., C.W., D.S., S.J., and H.R. are full-time employees at Biogen, Inc. The funders had no role in study design, data collection and analysis, decision to publish, or preparation of the manuscript.

Further information and requests for resources and reagents should be directed to and will be fulfilled by the Lead Contact, Heiko Runz

## METHODS

### Gene Selection

We chose to study 30 candidate genes from 18 loci reported as associated through common-variant genome-wide association studies (GWAS) as associated with plasma lipid levels and the risk for CAD. Twenty-eight of these genes had been identified and validated as functional regulators of LDL-uptake and/or cholesterol levels into cells in a previous RNAi-screen analysing a total of 133 genes in 56 lipid and CAD GWAS loci^17^ (**Table S1**). Common-variant association signals and published biological evidence for potential roles in lipid regulation were updated for all 30 candidate genes based on the recent literature (e.g., ^1–3,49^) and queries using the PhenoScanner platform^50^ (http://www.phenoscanner.medschl.cam.ac.uk/). Twenty eight genes were validated to reside within loci that are associated at genome-wide significance (p<5e-8) with plasma lipid levels or CAD. SNPs near *FAM174A* (rs383830) and *SEZ6L* (rs688034) had originally been reported as associated with CAD^51^, but failed to replicate at genome-wide significance in more recent meta-GWAS. However, since knockdown of both genes had scored as significantly impacting lipid parameters in cells^17^ the two genes were maintained for this current study.

### Colocalization Analysis

Colocalization analysis was performed between the 28 GWAS lead SNPs using summary statistics from the 2013 Global Lipid Genetics Consortium GWAS^1^ (http://csg.sph.umich.edu/willer/public/lipids2013/) and the GTEx liver *cis*-eQTL dataset (N=153)^52^. When a respective locus was associated with multiple lipid phenotypes, the SNP with the lowest reported p-value association with LDL was chosen to be the lead SNP. There was no GTEx liver expression data for four genes (*APOE, MYBPHL, NCAN, SEZ6L*), therefore there were no *cis*-eQTL for these genes to colocalize with. Colocalization analysis was conducted following the methods in Giambartolomei et al., 2014^53^ using the R ‘coloc’ package on a +/-500kb window around each lead SNP against SNP-to-expression data of all neighbouring genes within that locus. Positive colocalization between liver *cis*-eQTL and GWAS signal was defined as showing a posterior probability of sharing the same SNP (PP4) if larger than 0.8. A lead SNP at the *SORT1*/*CELSR2* locus (rs629301) showed a positive colocalization signal, but the *cis*-eQTL co-localized with both genes, so SNP-based GIs for these genes could not be analysed separately.

### UK Biobank lipid and CAD phenotypes

The UK Biobank is a prospective study of over 500,000 participants recruited at an age of 40-69 years from 2006-2010 in the United Kingdom. Participant data include health records, medication history and self-reported survey information, together with imputed genome-wide genotypes and biochemical measures^20^. Baseline biochemical measures including LDL cholesterol (LDL), HDL cholesterol (HDL), triglycerides (TG), and serum total cholesterol (TC) had been obtained in UK Biobank’s purpose-built facility in Stockport as described in the UK Biobank online data showcase and protocol (www.ukbiobank.ac.uk). Demographic and other relevant phenotypic information was obtained from standard questionnaire data. Individual lipid phenotypes (LDL, HDL, TG and TC) were modelled as dependent variables using linear regression models against covariates including age, sex, smoking, alcohol drinking status, and BMI. Lipid medication use was obtained from self-reported questionnaire data (UK Biobank fields 6153 and 6177). CAD cases were recognized based on both self-reported diagnosis and Hospital Episode Statistics data in the UK Biobank with a code-based CAD definition as presented in the most recent CAD GWAS that included UK Biobank^49^. In total, 30,125 CAD cases were identified and the cohort was adjusted for age, sex, smoking status, alcohol drinking status, BMI and lipid medication use. All phenotype data were derived from UK Biobank basket “ukb27390” released on March 11, 2019.

### Pairwise gene-based PTV-burden interaction testing

High-impact protein-truncating variants (PTVs) expected to disrupt protein functions were identified from 200,654 whole-exome sequencing (WES) data of UK Biobank participants to conduct pairwise interaction analyses. WES data was generated and quality controlled (QC-ed) as described in Van Hout et al. at the Regeneron Genetics Center as part of a collaboration between AbbVie, Alnylam Pharmaceuticals, AstraZeneca, Biogen, Bristol-Myers Squibb, Pfizer, Regeneron and Takeda and the UK Biobank consortium^54^. PTVs were called from a Regeneron QC-passing “Goldilocks” set of genetic variants using Variant Effect Predictor v96^23^ (McLaren et al., 2016) and the LOFTEE plugin^21^. We identified 462,762 high-confidence PTVs with a minor allele frequency of <1% in the canonical transcripts of 18,869 genes. This set included 755 rare PTVs in the 30 lipid genes analysed in this study. PTVs per gene were enumerated, and a PTV-burden association analysis was conducted in 161,508 unrelated (>2^nd^ degree relatedness) UK Biobank participants of European ancestry, as defined by principle components analysis of the genotyping data^20^. Replication analysis was conducted from an additional 101,827 samples, bringing the total sample size used for calling PTVs from UK Biobank exome sequencing data to 302,634. Of these 101,827 samples, 79,462 fulfilled the criteria applied to the discovery cohort, so that an overall sample size of 240,970 exomes was available for replicating findings from the initial PTV-based GI analyses.

For pairwise PTV-based interaction testing, QC-ed UK Biobank lipid phenotypes (HDL, LDL, TG and TC) were modelled as dependent variables using the following four linear regression models in R:

Model 1 for gene1 PTV-burden only: lipids ~ PTV_1_
Model 2 for gene2 PTV-burden only: lipids ~ PTV_2_
Model 3 for gene1 PTV-burden and gene2 PTV burden (*additive GI*): lipids ~ PTV_1_ + PTV_2_
Model 4 for gene1 PTV-burden and gene2 PTV burden (*non-additive GI*): lipids ~ PTV_1_ + PTV_2_ + PTV_1_ * PTV_2_

Schwarz’s Bayesian Information Criterion (BIC)^55^ scoring was used to determine the best model to explain the data and goodness of fit, with the lowest BIC value indicating the best-fitting model describing each gene pair. Model 3 reflected *additive* genetic interactions (aGIs), Model 4 *non-additive* gene interactions (naGIs). The model with the lowest BIC was chosen as describing most adequately the type of interaction between each corresponding gene pair.

### Pairwise SNP interaction testing

To assess whether GWAS lead SNPs modulate plasma lipid levels through joint effects within and across GWAS loci, we conducted pairwise SNP-SNP interaction analysis using genome-wide genotyping data and biochemical measures of lipid species from the UK Biobank. Twenty-eight lead SNPs mapped to the 30 lipid GWAS genes were extracted from genotyping data of 378,033 unrelated (removed up to 2^nd^ degree relatedness) participants of European ancestry. A total of 378 pairwise modifier effects were tested by conducting Robust Linear Model Fitting using R, running the following four linear regression models:

Model 1 for SNP1 only: lipids ~ SNP_1_
Model 2 for SNP2 only: lipids ~ SNP_2_
Model 3 for SNP1 and SNP2 (*additive GI*): lipids ~ SNP_1_ + SNP_2_
Model 4 for SNP1 and SNP2 (*non-additive GI*): lipids ~ SNP_1_ + SNP_2_ + SNP_1_ * SNP_2_

Schwarz’s Bayesian Information Criterion (BIC) scoring was used to determine the best model to explain the data and goodness of fit, with the lowest BIC value indicating the best-fitting model describing each SNP pair. If Model 3 had the lowest BIC value, it reflected an aGI, and if Model 4 had the lowest BIC value, it reflected a naGI.

A similar strategy was applied for pair-wise interaction testing to explore potential joint effects between the 30 genes on CAD risk by running the following four logistic regression models adjusted for age, sex, smoking status, alcohol drinking status, BMI and lipid medication use:

Model 1 for SNP1 only: CAD ~ SNP_1_
Model 2 for SNP2 only: CAD ~ SNP_2_
Model 3 for SNP1 and SNP2 (*additive GI*): CAD ~ SNP_1_ + SNP_2_
Model 4 for SNP1 and SNP2 (*non-additive GI*): CAD ~ SNP_1_ + SNP_2_ + SNP_1_ * SNP_2_

As above, the model with the lowest BIC was chosen as describing most adequately the type of interaction between each corresponding SNP pair.

### PTV-SNP interaction testing

In order to conduct pairwise interaction analyses between GWAS lead SNPs and PTVs, we assessed the interaction of the 28 lead SNPs with rare PTV burden for each of the 30 genes. For SNP-PTV interaction testing, UK Biobank lipid phenotypes (HDL, LDL, TG and TC) were modelled as dependent variables using the following four linear regression models:

Model 1 for gene1 lead SNP only: lipids ~ SNP_1_
Model 2 for gene2 PTV-burden only: lipids ~ PTV_2_
Model 3 for gene1 lead SNP and gene2 PTV burden (*additive GI*): lipids ~ SNP_1_ + PTV_2_
Model 4 for gene1 lead SNP and gene2 PTV burden (*non-additive GI*): lipids ~ SNP_1_ + PTV_2_ + SNP_1_*PTV_2_

As above, the model with the lowest BIC was chosen as describing most adequately the type of interaction between each corresponding SNP-gene pair.

### PTV-PRS interaction testing

We assessed the interaction effects between polygenic risk score (PRS) and PTVs for each of the four lipid phenotypes. To construct PRS for UK Biobank samples, we first derived the PRS weights for each SNP across the genome using PRS-CS^26^, which is a Bayesian regression-based algorithm, and publicly available summary statistics from lipid GWAS^1^. We applied derived PRS weights to imputed genotypes (with minor allele frequency >0.01 and imputation quality INFO >0.8) of UK Biobank samples and calculated PRS for each lipid, based on the corresponding PRS weights. Note that all SNPs in the gene of interest were excluded from the PRS when testing for PRS-PTV gene interaction. GIs were tested between PRS and PTV-burden for each of the 30 genes by fitting the four linear regression models:

Model 1 for PRS only: lipids ~ PRS
Model 2 for gene PTV-burden only: lipids ~ PTV
Model 3 for PRS and gene PTV burden (*additive GI*): lipids ~ PRS + PTV
Model 4 for PRS and gene PTV burden (*non-additive GI*): lipids ~ PRS + PTV + PRS * PTV

As above, the model with the lowest BIC was chosen as describing most adequately the type of interaction between each corresponding PRS-gene pair.

### RNAi interaction testing

#### Cells and reagents

HeLa-Kyoto cells are a strongly adherent Hela isolate (gift from S. Narumiya, Kyoto University Japan) that, as we demonstrated earlier, enable reliable measurements of LDL-cholesterol uptake dynamics and show lipid homeostatic mechanisms similar to those described for liver-derived cell models^17,27,28^. DiI-LDL (Life Technologies), DRAQ5 (Biostatus), Dapi (Molecular Probes), 2-hydroxy-propyl-beta-cyclodextrin (HPCD) (Sigma), Lipofectamine 2000 (Invitrogen) and Benzonase (Novagen) were purchased from the respective suppliers.

#### siRNA selection and production of siRNA microarrays

RNA-interference (RNAi) screening was conducted in glass-bottomed single-well chambered cell culture (Lab-Tek) slides with solid-phase reverse siRNA-transfection of cultured cells (“cell microarrays”) as described previously^27,56^. Each gene under study was targeted with a single siRNA (Silencer Select, Invitrogen) that had been selected with the EMBL-generated software tool bluegecko (J.K. Hériche, unpublished) based on the alignment to the reference genome, a maximal number of protein-coding transcripts per gene targeted and expected specificity for the target gene. The 28 siRNAs in this study had been validated earlier to significantly enhance or reduce cellular uptake of fluorescent-labelled LDL (DiI-LDL) or free cellular cholesterol levels^17^ and were shown to efficiently downregulate mRNA or protein levels of their respective target genes (**Table S2**). siRNA sequences are provided in Blattmann et al., 2013 Supplementary Table 4. For the two genes not analysed in our earlier study (*MYLIP*, *PAFAH1B2*), siRNAs used in the current study were prioritized from 3 and 5 siRNAs per gene based on bluegecko *in silico* analyses, knockdown efficiency on target mRNA/protein levels (up to less than 10% residual levels) and/or efficiency to modulate cellular DiI-LDL uptake in preparatory individual single gene knock-down experiments (not shown). The 75% (12/16) of siRNAs that had scored as individually modulating cellular DiI-LDL uptake in our earlier study^17^ also met the more stringent criteria of our current study to score as LDL-uptake modulator when used either alone or together with non-silencing control siRNA Neg9 (**Figure 3b**, CTRL column), thereby replicating our earlier results and validating experimental settings for this current study.

To cover the total of 435 pairwise siRNA combinations including controls and replicas, five different cell microarrays with 384 spots/array were produced. Per array, the following negative controls were added: eight spots containing *INCENP*-siRNA (s7424) to control for transfection efficiency^17^; eight spots containing non-silencing control siRNA Neg1 (s229174), and eight spots containing non-silencing control siRNA (denoted as CTRL throughout the text) Neg9 (s444246). Furthermore, eight spots were added with siRNA targeting *LDLR* (s224006) as a positive control for LDL uptake, as well as eight spots with siRNA targeting *NPC1* (s237198) knockdown of which increases free cellular cholesterol signals^27^. For pairwise combinatorial RNAi-screening, siRNAs against two genes were printed simultaneously on a respective siRNA-spot, with equal amounts (15 pmol/siRNA) of siRNA per gene. As positive controls, eight spots containing both, non-silencing control siRNA Neg9 (CTRL) (s444246) and siRNA targeting *LDLR* (s224006), and eight spots containing both, non-silencing control siRNA (CTRL) Neg9 (s444246) and siRNA targeting *NPC1* (s237197) were included per array. For all genes, “single-gene knockdown” scenarios [siRNA_*geneA*_+Neg9] were added on two spots per array. Each pairwise “combinatorial knockdown” scenario [siRNA_*geneA*_+siRNA_*geneB*_] was analyzed on one spot per array, with a single spot covering 50-100 informative cells^28,57^ (**Figure S1**).

In order to confirm GIs identified with the coRNAi screen, we replicated our analyses with forward transfection in a liquid-phase format with Lipofectamine 2000 reagent in 12-well plates, according to the manufacturer’s instructions. Concentrations of the siRNAs were adjusted to mimic the single knockdown phenotypes from the screen (**Table S2)**. 1μl of Lipofectamine 2000 was used per each transfection. GIs that showed a statistically significant interaction effects (p_fdr_<10^−2^) in replication analyses and acted in the same direction (same directionality of interaction value) as in the coRNAi screen, were considered as validated (**Table S12**).

#### Cell culture, transfection and LDL-uptake assay

HeLa Kyoto cells were grown in DMEM medium (Gibco) supplemented with 10 % (w/v) fetal calf serum (FCS)(PAA) and 2 mM L-glutamine (Sigma) at 37 °C with 5 % CO_2_ and saturated humidity. Cells were plated at a density of 6×10^4^ per plate on the cell microarrays for solid-phase siRNA transfection^56^ and cultivated for 48 hours before performing the LDL-uptake assay. For liquid phase transfection-based validation experiments, cells were plated in 12-well plates the day prior to transfection, and siRNA-transfected cells were cultivated for 48 hours. The assays to monitor cellular uptake of fluorescently-labelled LDL (DiI-LDL) were performed as described in more detail in previous publications^17,27,28^. In brief, cells cultured in serum-free medium (DMEM/2mM L-glutamine/0.2 % (w/v) BSA) and exposed to 1% 2-hydroxy-propyl-beta-cyclodextrin for 45min were labelled with 50 μg/ml DiI-LDL (Invitrogen) for 30 min at 4 °C. DiI-LDL uptake was stimulated for 20 min at 37.0 °C before washing off non-internalized dye for 1 min in acidic (pH 3.5) medium at 4 °C, fixation, and counterstaining for nuclei (Dapi) and cell outlines (DRAQ5). For RNAi-based gene interaction screening, each of the five cell microarrays was assayed in 7-10 biological replicas.

#### Image acquisition and quality control

Image acquisition was performed using an Olympus IX81 automated microscope with Scan^R software and an UPlanSApo 20x/NA 0.40 air objective as described^17,28^. Images from a total of 42 cell microarrays were visually quality controlled. Arrays with insufficient knockdown efficiency where *INCENP* siRNA treated cells did not show the expected multinucleated phenotype in the DAPI channel were excluded. Also arrays with plate effects as evaluated through diagnostic plots with the *splot* function in R, and arrays where knockdown of *LDLR*, or *LDLR* together with negative control siRNA Neg9, did not show a significant difference from controls, were discarded as well. Following these QC criteria, 29 cell microarrays with a total of 11,047 image frames per channel were further analysed. The in-house developed tool HTM Explorer (Ch. Tischer; https://github.com/embl-cba/shinyHTM) was then used to select images fulfilling pre-defined criteria for cell number, image sharpness quality, and image background intensity, resulting in a total number of 9,539 (86.35%) QC-ed image frames that were used for subsequent analyses.

#### Image analysis

Automated image analysis was performed using a specifically developed pipeline (available upon request) in the open source software CellProfiler^58^ http://www.cellprofiler.org as described^17,28^. In brief, areas of individual cells were approximated by stepwise dilation of masks on the DAPI (nuclei) and DRAQ5 (cell outlines) channels^59^. For each individual cell, DiI-LDL signal was determined from masks representing intracellular endosome-like vesicular areas that were determined by local adaptive thresholding according to predefined criteria for size and shape (**Figure S1**). Total fluorescence intensity of DiI-signal above local background per cell mask was quantified, and means were calculated from all cells per image. Then, for each siRNA, or siRNA combination (“*treated*”), signals from different images from the same biological replicate were averaged and a robust Z-score was calculated using the median fluorescence signal of all the negative control siRNAs per array (“*median(controls)*”) and by the median absolute deviation of these controls (“mad(controls)”) as follows: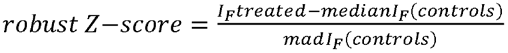^60,61^A median robust Z-score was calculated per treatment across all biological replicates and is represented on the plots.

#### RNAi gene interaction testing

To identify pairs of genes for which simultaneous knock-down results in an additive or non-additive gene interaction effects on LDL uptake we conducted a Robust Linear Model fitting in R. RobustZScore values calculated from different biological replicates in the presence of single ([siRNA_geneA_+Neg9] and [siRNA_geneB_+Neg9]) or double knock-downs ([siRNA_geneA_+ siRNA_geneB_]) were considered to be response variable value. Negative control values [Neg9] were included in each fitted dataset to correctly account for baseline LDL uptake. The full regression model considered in the study was

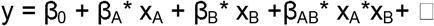

which is equivalent to the short form of the statistical formula:

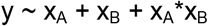

In both formulas y corresponds to the robustZscore values of measured LDL uptake; x_A_, x_B_ are encoded predictor variables, which are equal to 1 in case of presence of siRNA_geneA_, siRNA_geneB_, or both siRNAs accordingly and equal 0 otherwise. The ◻ is a noise term, which is minimised during the fitting process. Model fitting provides estimates of β_0_, β_A_, β_B_ and β_AB_ values. β_0_ defines the effect of the negative control on robustZscore values and can be also denoted as an intercept of the linear fit. For our data β_0_ is always close to 0 because of the robustZscore definition. The β_A_ and β_B_ define individual effects of siRNA_geneA_ and siRNA_geneB_ accordingly. The β_AB_ defines the interaction effect between genes A and B and represents the difference between the observed robustZscore values in case of double knockdown y_AB_ and the expected additive effect of geneA and geneB knockdown (β_AB_ = y_AB_ - β_0_ - β_A_ - β_B_).

Subsequently, two strategies were used to evaluate functional interactions for each gene pair using defined statistical model:

First, to determine gene pairs for which genetic interactions and additive effects observed upon combinatorial knockdowns, we compared fitting of the whole model to the fitting of reduced model versions. Following models were compared:

Model 0 - (only baseline effect β_0_ in case of either single or double knockdown): y ~ 1
Model 1 - effect of siRNA_geneA_ only: y ~ x_A_
Model 2 - effect of siRNA_geneB_ only: y ~ x_B_
Model 3 for additive effect of both siRNAs (additive GI): y ~ x_A_ + x_B_
Model 4 – full model including genetic interaction (non-additive GI): y ~ x_A_ + x_B_ + x_A_*x_B_.

To determine the best model explaining the data for each gene pair we used Schwarz’s Bayesian Information Criterion (BIC). BIC score was calculated for each model fitted to the data, then the model with the lowest BIC value (BIC*) was selected as the best-fitting model. Co-knockdown effects of each gene pair were classified as aGIs or naGIs when model 3 or model 4 accordingly were defined to fit data best. Additionally, for the RNAi screen, we used the method published by Raftery, 1995 to define the strength of evidence for the respective model to be selected^62^. Namely, if the difference (ΔBIC) between the BIC value of the best fitting model (the model with the lowest BIC value) and the BIC value of any other model is bigger than 2, then it would indicate a significant evidence for this model (with BIC*) to truly represent the data. In other words, if ΔBIC>2 then the model with lowest BIC value (BIC*) was considered as the one most correctly describing the data in comparison to other tested models. If the ΔBIC<2, then two models were considered as possible alternatives for representing the dataset.

Secondly, to estimate statistical significance of gene interaction effect for each siRNA gene combination, we calculated a p-value from the t-value of the linear regression model term, describing genetic interaction (β_AB_) as p_Val_=2-2*p_norm_(abs(t_Val_)). To correct for multiple comparisons, the p-values were adjusted using the false discovery rate (fdr) method^63^, and the fdr-corrected p-values < 10^−2^ were considered to correspond to significant GIs.

#### RT-qPCR analysis

Cell lysis and total RNA extraction was done using the RNease Mini Kit (Qiagen). Reverse-transcription was performed with the SuperScript™ III First-Strand Synthesis SuperMix for RT-qPCR (Invitrogen). RT-qPCR data was obtained from three biological replicates/siRNA treatment. Primers can be provided upon request. For each siRNA treatment target mRNA was normalized to that of *GAPDH* and compared to CTRL siRNA and the log2 of fold change (2^−^ΔΔCT) was calculated (see **Figure S4**). Results were considered as significant if p-values were below 0.05 in a two-tailed Student’s t-test.

#### Immunocytochemistry and confocal microscopy

Cells were transfected and cultured as described above then fixed with 3% paraformaldehyde (PFA) at room temperature for 20 min, washed with PBS, incubated with 30mM glycine for 5 min and washed again with PBS. For LDLR staining cells were permeabilized with 0.05% Filipin III (Sigma #F4767) in 10% FCS for 30 min at room temperature. Primary antibody: rabbit monoclonal anti-LDLR (Fitzgerald #20R-LR002) was diluted in 5% FCS overnight at 4 °C. Secondary antibody: goat polyclonal goat anti-rabbit IgG Alexa 568 (Invitrogen #A11011) was diluted 1:400 in 5% FCS. Fixed cells were imaged using a Zeiss LSM 780 confocal microscope using a 63x/NA 1.4 oil objective.

