## Supplementary Figures for "Pairwise genetic interactions modulate lipid plasma levels and cellular uptake": Zimon_SupplementaryFigures.pdf

**Contents**

Supplementary Fig. 1: Cellprofiler pipeline for quantitative image analysis of LDL-uptake assay
Supplementary Fig. 2: Boxplot representation of the 20 gene-gene interactions that were replicated with liquid phase transfection.
Supplementary Fig. 3: Subcellular localization of LDLR upon knockdown of interacting genes.
Supplementary Fig. 4: No effect of *TOMM40* knockdown on *APOE* expression levels.

### Supplementary Tables

Supplementary Table 1: GWAS support for 30 lipid/CAD candidate genes analyzed in this study
Supplementary Table 2: *A priori* evidence for lipid/CAD-relevant biological functions of the candidate lipid genes under study
Supplementary Table 3: PTVs identified in the 30 lipid/CAD GWAS genes through exome sequencing of 200,654 UK Biobank participants
Supplementary Table 4: Single-gene PTV-burden association results for the 30 lipid/CAD GWAS genes with four lipid traits
Supplementary Table 5: Pairwise gene-based PTV-PTV burden interaction analysis results in 161,508 UK Biobank participants
Supplementary Table 6: Replication analysis of PTV-based GIs in an additional 79,462 UK Biobank participants
Supplementary Table 7: Pairwise lipid/CAD GWAS lead SNP-SNP interaction analysis results in the UK Biobank for the 28 loci analyzed in this study
Supplementary Table 8: Results for testable pairwise GWAS lead SNP-PTV burden interaction analysis
Supplementary Table 9: Results for modifier effects between PRS and PTV burden interaction analysis
Supplementary Table 10: Results of primary coRNAi screen analyzed via Bayesian Information Criterion (BIC) model fitting
Supplementary Table 11: Results of primary coRNAi screen analyzed via robust linear model fitting
Supplementary Table 12: Replication of coRNAi screening results independently validate 20 gene pairs as showing non-additive effects on cellular LDL-uptake
Supplementary Table 13: Comparative analysis of additive effects identified gene pairs by RNAi screen

**Figure S1**

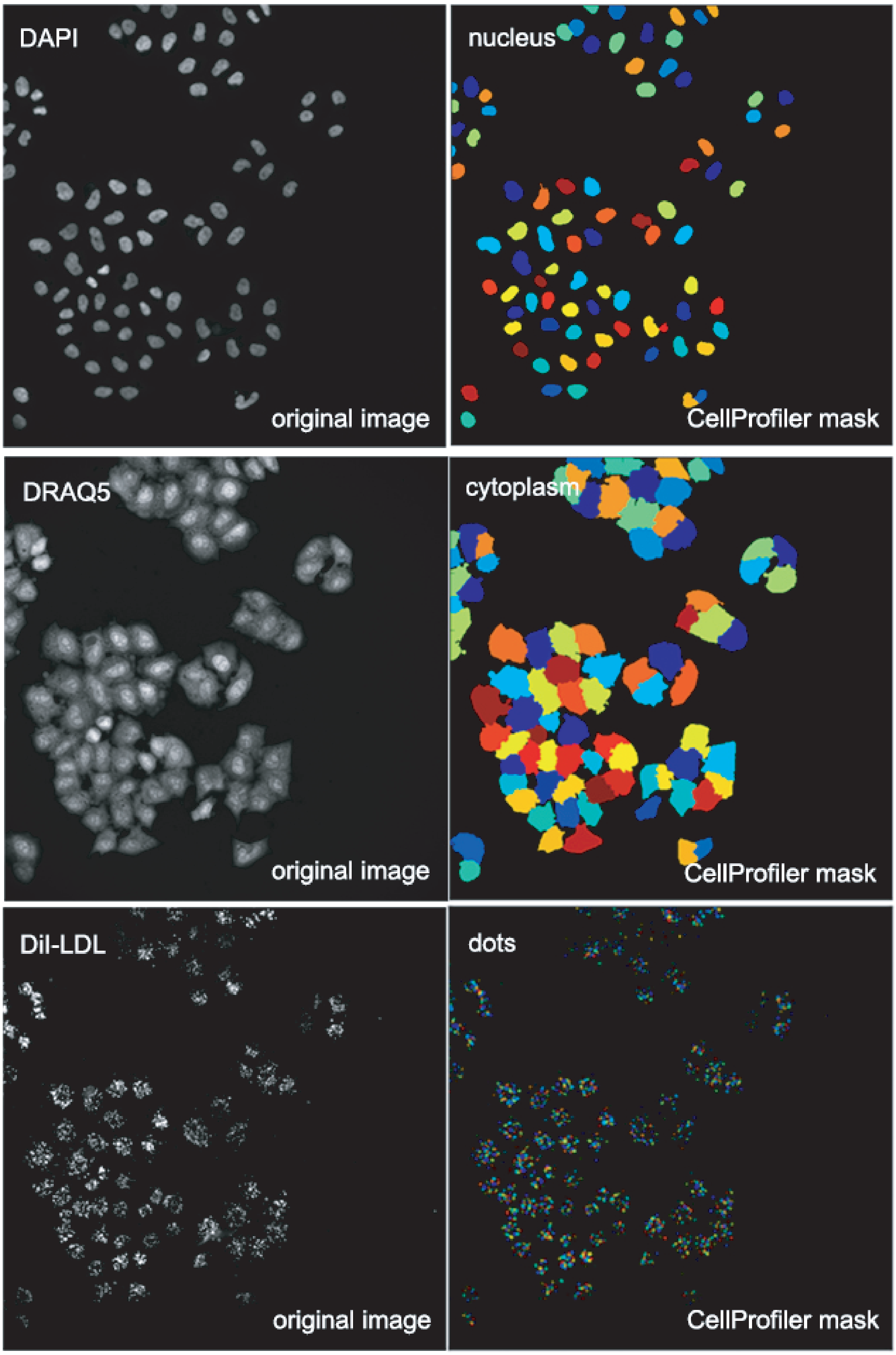


**Figure S1. Cellprofiler pipeline for quantitative image analysis of LDL-uptake assay.** Shown are representative images from the LDL-uptake assay (left column) acquired with the Scan^R software of the Olympus widefield microscope with the 20x objective and the segmentation of cellular structures (right column) performed through a Cellprofiler pipeline.

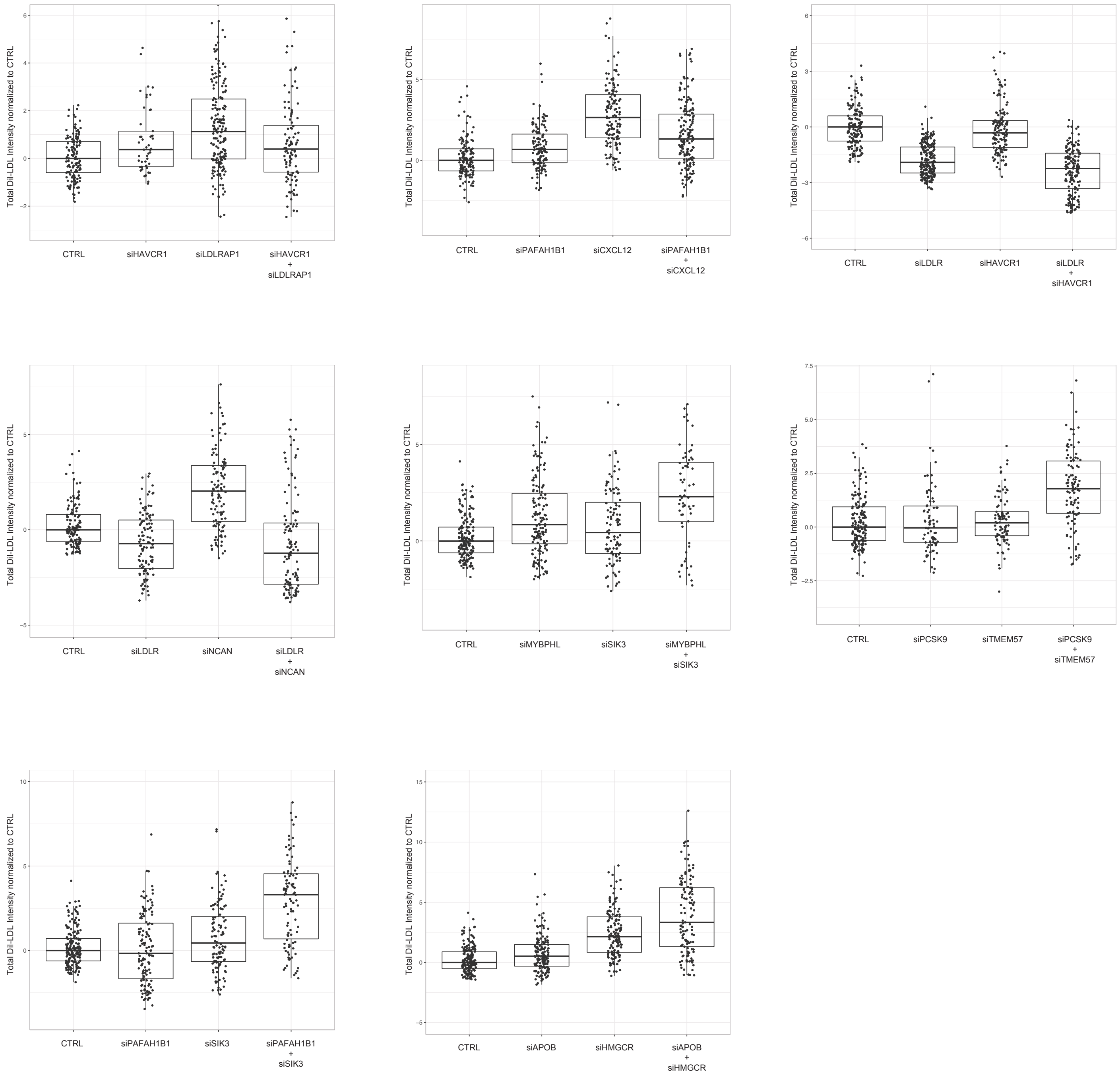


**Figure S2. Boxplot representation of the 20 gene-gene interactions that were replicated with liquid phase transfection.**

Shown are the 20 gene-gene interactions that were validated with liquid-phase transfection.

The boxplots show the median intensity of internalized DiI-LDL, normalized to the control, for the two single knockdowns (transfected together with the control siRNA), as well as for the double knockdown, for each pair of genes. (n=3-4)

**Figure S3**

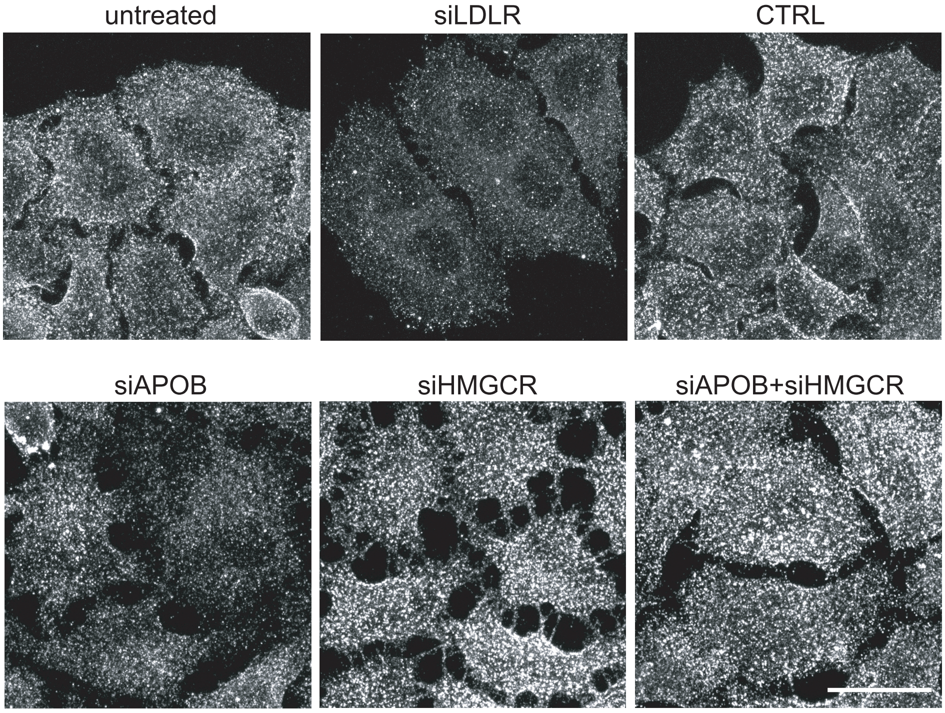


**Figure S3. Subcellular localization of LDLR upon knockdown of interacting genes.**
HeLa Kyoto cells were transfected with siRNAs targeting products of indicated genes and stained with antibody against LDLR. Cells were grown under sterol-depleted conditions (see Materials and Methods). Shown are maximal projections of confocal stacks of representative cells. Bar=20 μm.

**Figure S4**

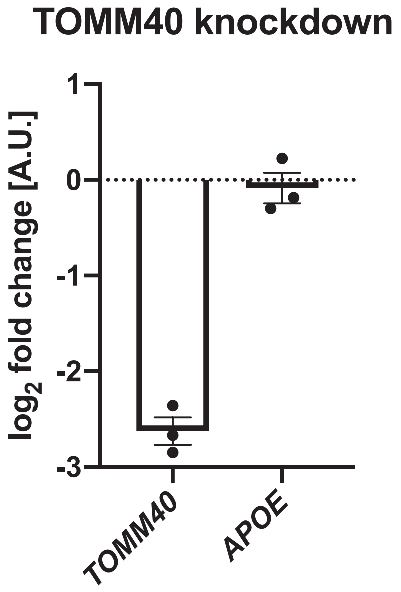


**Figure S4. No effect of *TOMM40* knockdown on *APOE* expression levels.**
Shown are mRNA levels of either *TOMM40* or *APOE* in HeLa Kyoto cells after 48 h knockdown of *TOMM40* after normalization to the control siRNA. The target gene mRNA levels were normalized to the housekeeping gene, *GAPDH*. The error bars represent the standard error of the mean (n=3).
